# A modular cloning toolkit for genome editing in plants

**DOI:** 10.1101/738021

**Authors:** Florian Hahn, Andrey Korolev, Laura Sanjurjo Loures, Vladimir Nekrasov

## Abstract

**Background:** CRISPR/Cas has recently become a widely used genome editing tool in various organisms, including plants. Applying CRISPR/Cas often requires delivering multiple expression units into plant and hence there is a need for a quick and easy cloning procedure. The modular cloning (MoClo), based on the Golden Gate (GG) method, has enabled development of cloning systems with standardised genetic parts, e.g. promoters, coding sequences or terminators, that can be easily interchanged and assembled into expression units, which in their own turn can be further assembled into higher order multigene constructs.

**Results:** Here we present an expanded cloning toolkit that contains ninety-nine modules encoding a variety of CRISPR/Cas-based nucleases and their corresponding guide RNA backbones. Among other components, the toolkit includes a number of promoters that allow expression of CRISPR/Cas nucleases (or any other coding sequences) and their guide RNAs in monocots and dicots. As part of the toolkit, we present a set of modules that enable quick and facile assembly of tRNA-sgRNA polycistronic units without a PCR step involved. We also demonstrate that our tRNA-sgRNA system is functional in wheat protoplasts.

**Conclusions:** We believe the presented CRISPR/Cas toolkit is a great resource that will contribute towards wider adoption of the CRISPR/Cas genome editing technology and modular cloning by researchers across the plant science community.

## Background

The CRISPR/Cas technology has recently become an easily accessible genome editing tool for many organisms, including plants [1]. Generating gene knockouts has become a rather straightforward CRISPR/Cas application in many plant systems [2–4], while more sophisticated applications, such as allele replacements or targeted gene insertions, still remain a challenge due to low efficiency of homology-directed repair (HDR) in plants [5].

In its conventional form, the CRISPR/Cas system includes a DNA nuclease, such as Cas9, which is guided to a specific genomic location by the guide RNA. Therefore, in order to perform targeted mutagenesis *in planta*, one needs to co-express both the CRISPR/Cas nuclease and its cognate guide RNA. Usually, the gene encoding the CRISPR/Cas nuclease is expressed using an RNA polymerase II (Pol II) promoter (e.g. 35Sp), while the guide RNA is expressed under an RNA polymerase III (Pol III) promoter (e.g. U6p or U3p), which has a defined transcription start nucleotide (‘G’ for U6p or ‘A’ for U3p). One of the advantages of CRISPR/Cas is multiplexing i.e. one can target DNA at multiple genomic locations by co-expressing multiple guide RNAs specific to those loci. Guide RNAs can be expressed either as individual transcriptional units, each under its own Pol III promoter [4], or as a tRNA-sgRNA polycistronic transcript [6]. In the latter case, guide RNAs are interspaced with tRNAs in a single transcript that gets processed into individual guide RNAs by the highly conserved tRNA processing machinery inside the plant cell [6].

As genome editing applications in plants often rely on delivering multiple expression units into plant cells, including a selectable marker, a CRISPR/Cas nuclease-encoding gene and one or more guide RNAs, it is important to be able to assemble DNA constructs encoding such expression units easily and rapidly. The modular cloning (MoClo) system based on the Golden Gate (GG) cloning method [7] is highly flexible and versatile, and provides a means for quick and facile assembly of multi-expression unit constructs using standard genetic parts, such as promoters, terminators, coding sequences etc. The system has already been successfully used for genome editing applications in plants [3, 4, 8–10] but lacks modules encoding many of the newest genome editing reagents. Here we report on an expanded GG cloning toolkit for genome editing applications in monocot and dicot plants. We believe the toolkit will become a valuable addition to already existing GG-based tools for plant genome editing and be widely used by plant researchers across the community.

## Results

During this study we have generated a set of ninety-nine GG modules that enable one to perform genome editing in both monocot and dicot plant species (Additional file 2: Table S5). The cloning toolkit is an addition to previously published GG modules [7, 9–11] and includes new sequence-specific CRISPR/Cas nucleases (codon-optimised for both monocots and dicots), Pol II and Pol III promoters, as well as guide RNA backbone modules. The latter enable insertion of the guide sequence by cloning in a pair of annealed complimentary oligos without a PCR amplification step involved. The toolkit enables assembly of CRISPR/Cas constructs that target a single as well as multiple targets with guide RNAs expressed either under individual Pol III promoters (Fig. 1) or using a polycistronic tRNA-sgRNA construct (Fig. 4).

**Fig. 1.**
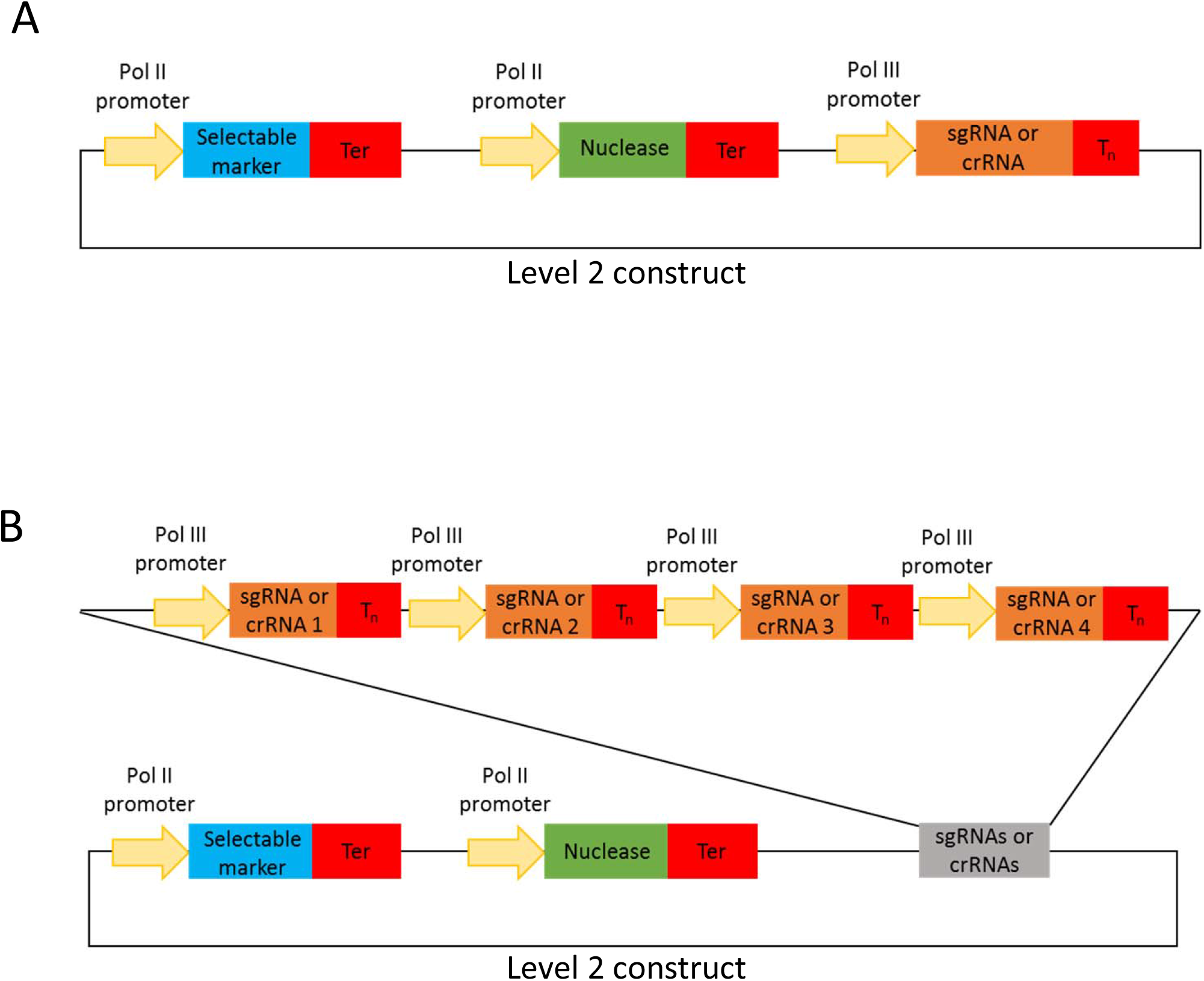
Cartoons of level 2 CRISPR/Cas constructs for targeted mutagenesis using guide RNAs expressed under individual Pol III promoters. (**a**) represents a construct with one and (**b**) – with four guide RNAs.

### CRISPR/Cas nuclease modules

The CRISPR/Cas nuclease level 0 modules include *Streptococcus pyogenes* Cas9 (SpCas9; pFH13, pFH24 and pFH25) as well as Cas9 variants coming from other bacterial species: *Staphylococcus aureus* (SaCas9; pFH14), *Streptococcus thermophilus* (StCas9; pFH15) and *Streptococcus canis* (ScCas9; pFH76) (Additional file 2: Table S5). SpCas9 [12] with the ‘NGG’ protospacer adjacent motif (PAM) is the most commonly used Cas9 variant for genome editing applications in various organisms, including plants. SaCas9 (‘NNGRRT’ PAM) and StCas9 (‘NNRGAA’ PAM) are less common but have also been successfully used in rice, tobacco [13] and Arabidopsis [14–16]. ScCas9 (pFH76), SpCas9-NG (pFH32) and SpCas9-derived xCas9 (pFH22) are characterised by broadened PAM motif requirements: ‘NNG’ for ScCas9 [17], and ‘NG’ for SpCas9-NG [18] and xCas9 [19]. We have also included modules with Cas12a (Cpf1) CRISPR nucleases from *Francisella novicida* (FnCas12a; pFH16 and pFH46) and *Lachnospiraceae bacterium* (LbCas12a; pFH17 and pFH47) as well as with four related Cms1 nucleases (pFH18-21) (Additional file 2: Table S5). LbCas12a [20], FnCas12a [21] and Cms1 [22] have all been shown to work in plants.

Base editors are a rather recent addition to the range of available genome editing tools and allow targeted conversion of DNA base pairs as following: C-G to T-A [23] and A-T to G-C [24] without introducing a double-strand break (DSB). The former base editor is based on the cytidine deaminase while the latter – on the adenosine deaminase. Both base editors have now been shown to be functional in various plants, including wheat, rice and tomato [25–29]. We have therefore generated level 0 modules encoding cytidine deaminase (pFH55 and pFH79) and adenosine deaminase (pFH45 and pFH92) based base editors (Additional file 2: Table S5).

EvolvR CRISPR-guided error-prone DNA polymerases have recently been shown to be able to introduce random point mutations at a targeted genomic locus [30]. Based on the Halperin et al. (2018) manuscript, we have generated a level 0 module with the wheat codon optimised version of enCas9–PolI3M–TBD (pFH77; Additional file 2: Table S5) that could prove to be a useful tool for reverse genetics in monocot plants.

We used the above mentioned CRISPR/Cas nuclease level 0 modules to assemble twenty-three nuclease expression units inserted into level 1 GG vectors to be applied in monocot and dicot plant species (Additional file 2: Table S5).

### Guide RNA modules

As CRISPR/Cas is an RNA-guided nuclease, guide RNA is its essential component that must be co-expressed with the nuclease in order to achieve on-target DNA cutting. Guide RNAs are usually expressed under Pol III promoters, such as U3p or U6p, that have a defined transcription start nucleotide (‘A’ and ‘G’, respectively). A number of genomic loci can be targeted simultaneously by CRISPR/Cas by co-expressing multiple guide RNAs and the modular cloning system is highly suitable for assembling constructs carrying multiple expression units, such as the CRISPR/Cas nuclease and guide RNAs.

As part of this study, we have generated a number of level 0 Pol III promoter modules (TaU3p, OsU3p, OsU6-2p and AtU6-26p; Additional file 2: Table S5). In addition, we have produced a number of guide RNA backbone level 0 constructs that can be used to assemble single or multiple guide RNA expression units without a PCR amplification step involved (Additional file 2: Table S5). The cloning system we present allows guide RNAs to be expressed either under individual Pol III promoters (Figs. 1–3; Additional file 1: section 2) or from a polycistronic tRNA-sgRNA construct, which includes sgRNAs interspaced with tRNA scaffolds [6] (Figs. 4 and 5; Additional file 1: section 3). The method with guide RNAs expressed under individual promoters enables expression of sgRNAs of the Cas9 family of CRISPR/Cas nucleases, which carry the guide sequence at the 5’ end of sgRNA (Fig. 2), as well as crRNAs of Cas12a (Cpf1) nucleases and related Cms1 nucleases, which have the guide sequence at the 3’ end of crRNA (Fig. 3).

**Fig. 2.**
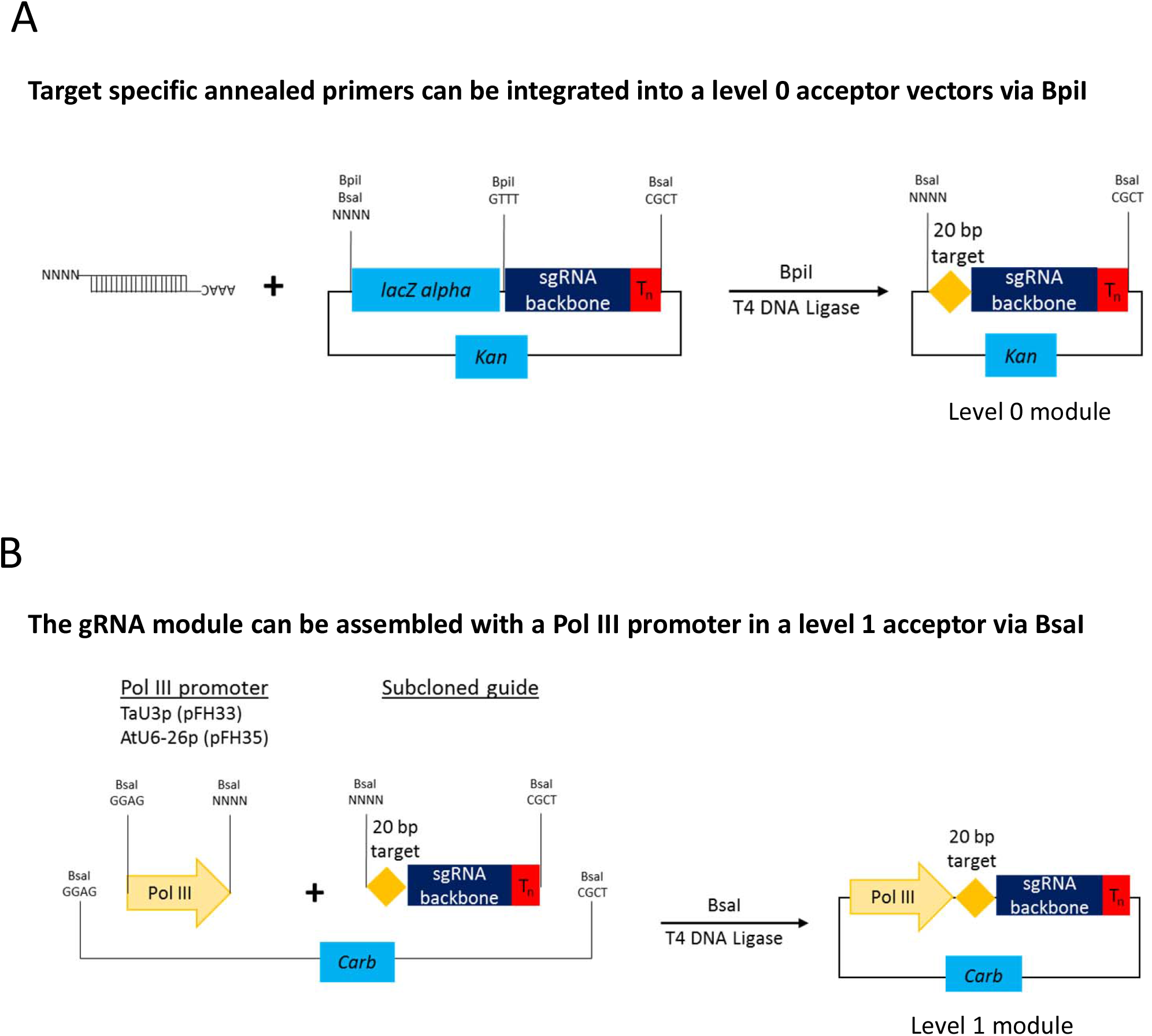
Assembly of level 1 sgRNA expression modules to be used with respective Cas9 nucleases. During the first step (**a**) annealed complementary oligos encoding the guide are inserted into a level 0 acceptor with the sgRNA backbone using BpiI. During the second step (**b**), sgRNA is fused with the respective Pol III promoter using BsaI.

**Fig. 3.**
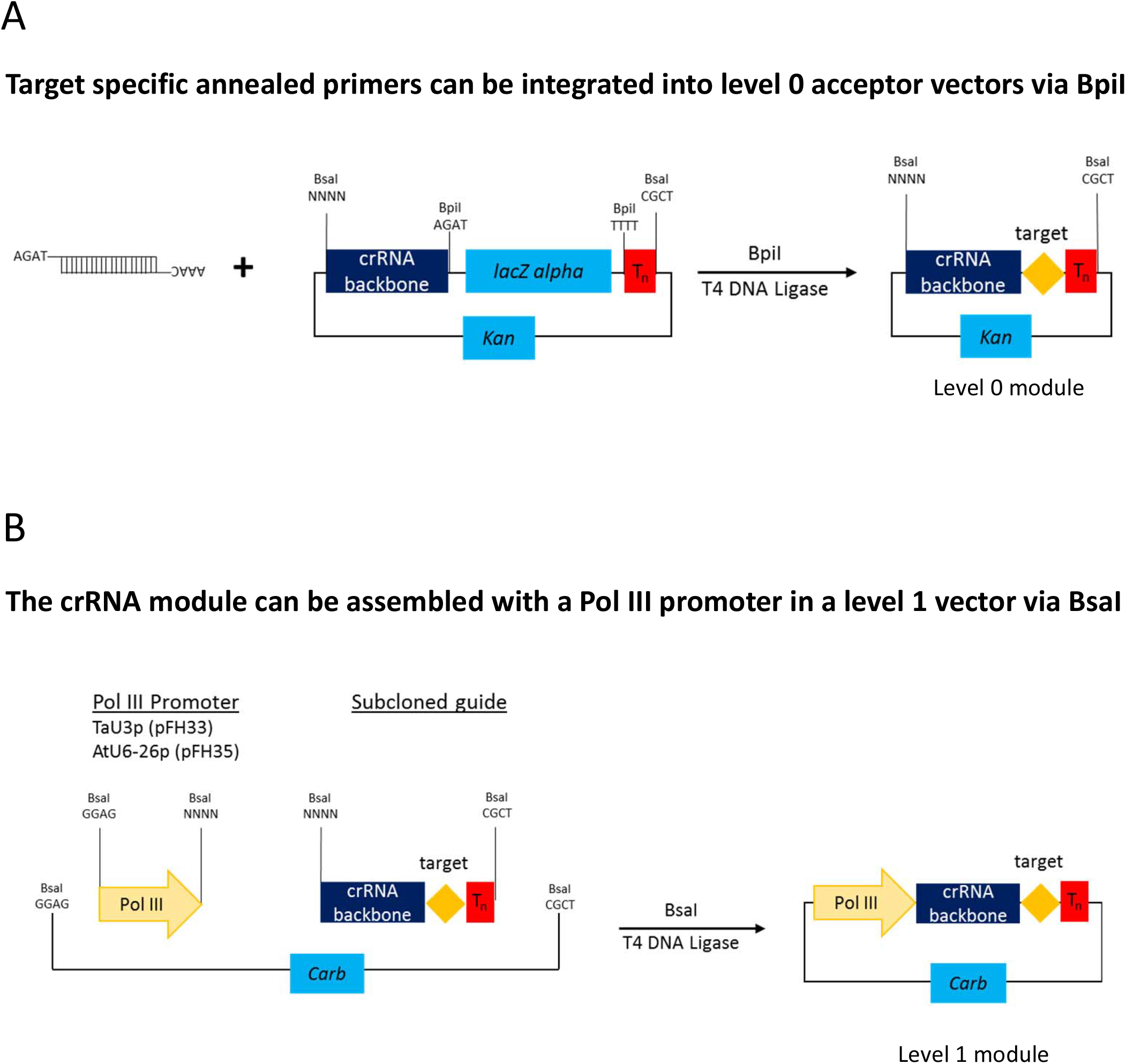
Assembly of level 1 crRNA expression modules to be used with respective Cas12a (Cpf1) and Cms1 nucleases. During the first step (**a**) annealed complementary oligos encoding the guide are inserted into a level 0 acceptor carrying the crRNA backbone using BpiI. During the second step (**b**), crRNA is fused with the respective Pol III promoter using BsaI.

Up to four guide RNAs under individual Pol III promoters can be assembled in using the former cloning procedure (Fig. 1b) and up to six sgRNAs per polycistronic construct – using the latter one (Fig. 4b). It must be noted that the number of guide RNAs under individual promoters could be increased to five, if no selectable marker is needed (Fig. 1b), or many more if level M/ level P vectors are used [11, 31]. As to tRNA-sgRNA polycistronic constructs, the total number of sgRNAs assembled into a level 2 destination vector could be up to twenty-four (six per level 1), with a selectable marker, and up to thirty, if no selectable marker is used. Again, it is possible to add more than thirty sgRNAs by using level M/ level P vectors [11, 31]. Our GG toolkit enables the user to build such complex constructs within a week (Additional file 1: Figure S1).

### Testing of the tRNA-sgRNA CRISPR/Cas constructs in wheat protoplasts

The GG toolkit therefore allows rapid parallel assembly of constructs by streamlining the cloning process. Since building multiple CRISPR constructs using GG is a straightforward procedure, it becomes reasonable to compare the activity of several experimental CRISPR setups in a transient expression system, such as protoplast, before proceeding with stable plant transformation, which could be a highly laborious and time consuming process. Our CRISPR toolkit includes three wheat codon optimised SpCas9 versions (level 0 constructs pFH13, pFH24 and pFH25; Additional file 2: Table S5) and their respective level 1 transcription units (pFH23, pFH66 and pFH67; Additional file 2: Table S5). These SpCas9 variants differ by e.g. nuclear localisation signal (NLS) versions or affinity tags. We have therefore compared the activity of the three Cas9 variants in wheat protoplasts by cotransforming each of the level 1 constructs (pFH23, pFH66 and pFH67) with the level 1 plasmid containing the six sgRNAs (Fig. 6a) assembled into a tRNA-sgRNA array. This has allowed us to target three different wheat genes at once (Fig. 6a). We have targeted each gene by at least two sgRNAs with large deletions between Cas9 cut sites expected to be detectable by PCR due to DNA band shifts as previously described [4]. PCR amplification of the target genes has revealed clear additional bands corresponding to alleles carrying large CRISPR/Cas-induced deletions in protoplasts transformed with pFH66. In contrast, application of the other two Cas9 versions (pFH23 and pFH67) resulted in very faint bands of the size corresponding to amplicons carrying the deletions (Fig. 6b). Our results therefore suggest a significantly higher activity of the pFH66-encoded SpCas9, as compared to the other two Cas9 variants, in wheat protoplasts.

## Discussion

The modular cloning kit presented in the study enables quick and facile assembly of DNA constructs for genome editing applications in plants and is an addition to previously published collections of compatible GG modules [7, 9–11]. The kit includes modules encoding a number of CRISPR/Cas nucleases (SaCas9, StCas9, LbCas12a etc.) that could be used as an alternative to the most commonly utilised SpCas9. SaCas9, for instance, has proven to be an efficient tool for generating gene knockouts in a number of plant species [13–16] and, in addition, has been shown to increase HDR efficiencies in plants [15]. Due to SaCas9 and StCas9 having longer than SpCas9 PAM motifs they are also likely to be more specific when it comes to DNA target recognition.

Cas12a (Cpf1) generates a staggered cut in DNA [32], while Cas9 – a blunt cut [12]. Due to this reason, Cas12a (Cpf1) could be a preferred choice of a CRISPR/Cas nuclease when it comes to HDR-based genome editing applications, such as targeted gene insertion [33, 34]. It is noteworthy that the modular cloning system is highly suitable for HDR-based applications as the DNA repair template could easily be cloned as a level 1 module into a level 2 destination vector. Also, since Cas12a (Cpf1) has a T-rich PAM motif (‘TTTN’ for LbCas12a (LbCpf1) and ‘TTN’ for FnCas12a (FnCpf1)), it could be a better choice for targeted mutagenesis in plant species with AT-rich genomes as compared to SpCas9 (‘NGG’ PAM).

As part of our study, we have generated a number of guide RNA backbone level 0 modules, which are compatible with respective Pol III promoters and CRISPR/Cas nucleases (Additional file 1: Table S1). The guide RNA backbone modules can be used for PCR-free assembly of guide RNA expression units by cloning in an annealed pair of complimentary oligos encoding the guide sequence (Figs. 2 and 3). These guide RNA backbones are to be used when one wishes to express guide RNAs under individual promoters and up to four guide RNAs can be assembled into a level 2 vector together with level 1 modules encoding a CRISPR/Cas nuclease and a selectable marker (e.g. BAR, NPTII etc.; Fig. 1, Additional file 2: Table S5 and Additional file 3: Table S6).

In addition to expressing each guide RNA under its own promoter, we have generated modules that allow assembly of polycistronic tRNA-sgRNA constructs with up to six sgRNAs expressed using a single Pol III promoter (Fig. 4). The tRNA-sgRNA system, originally described by Xie et al. (2015) in rice, was later successfully applied in wheat [35] and Arabidopsis [36].

**Fig. 4.**
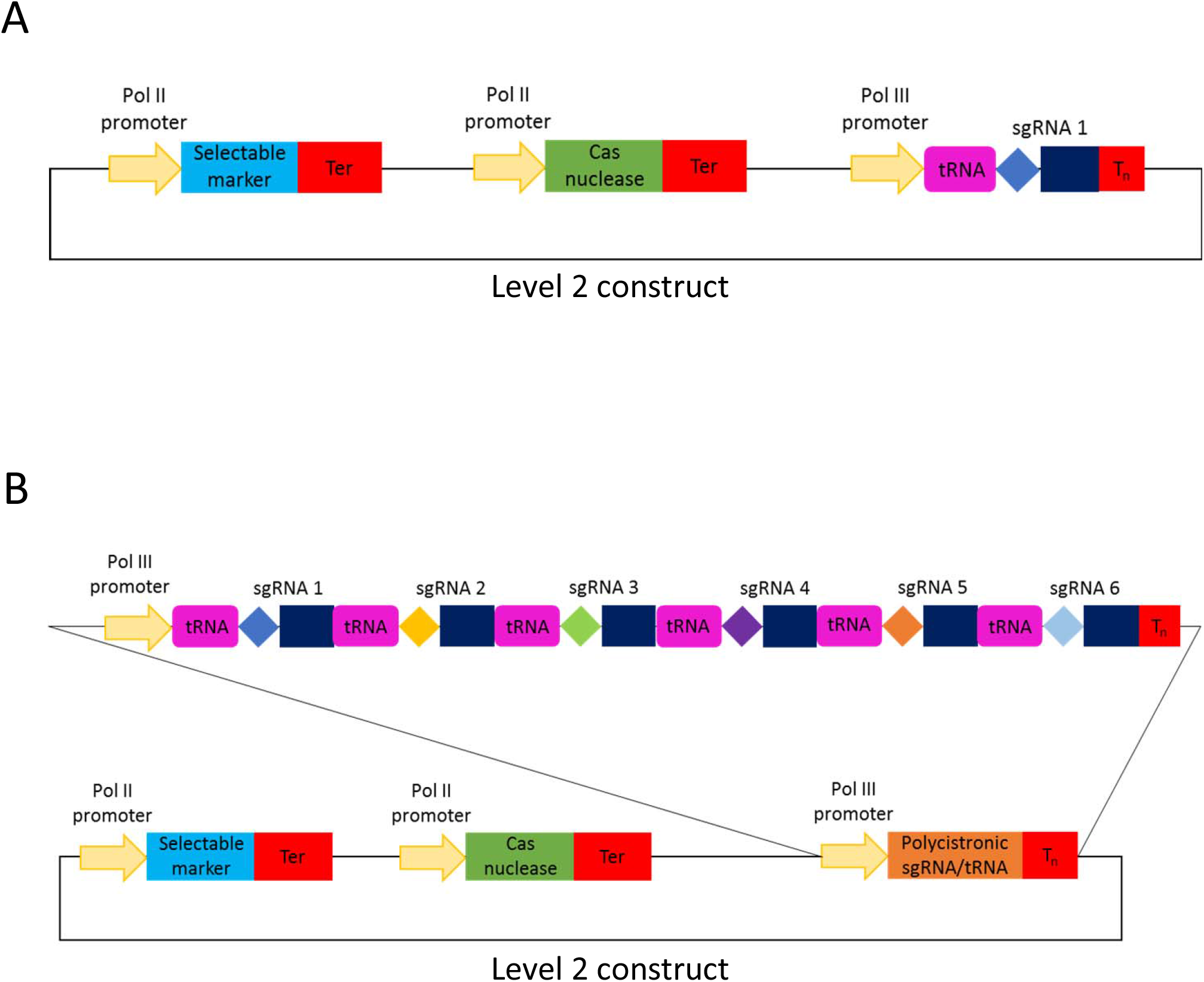
Cartoons of level 2 CRISPR/Cas constructs for targeted mutagenesis using the polycistronic SpCas9 tRNA-sgRNA system. (**a**) represents a construct with one and (**b**) – with six sgRNAs.

Nevertheless, the previously reported tRNA-sgRNA system relies on a rather cumbersome DNA construct assembly process as it involves PCR amplification of DNA fragments carrying repeats. The level 0 GG modules we have generated (Fig. 5 and Additional file 1: Table S3) enable straightforward and efficient assembly of tRNA-sgRNA arrays without a PCR step involved. The system offers a choice of five monocot and dicot Pol III promoters, with TaU6p being a published module [37], and two different SpCas9 sgRNA backbones (classic and improved; Additional file 1: Table S3).

**Fig. 5.**
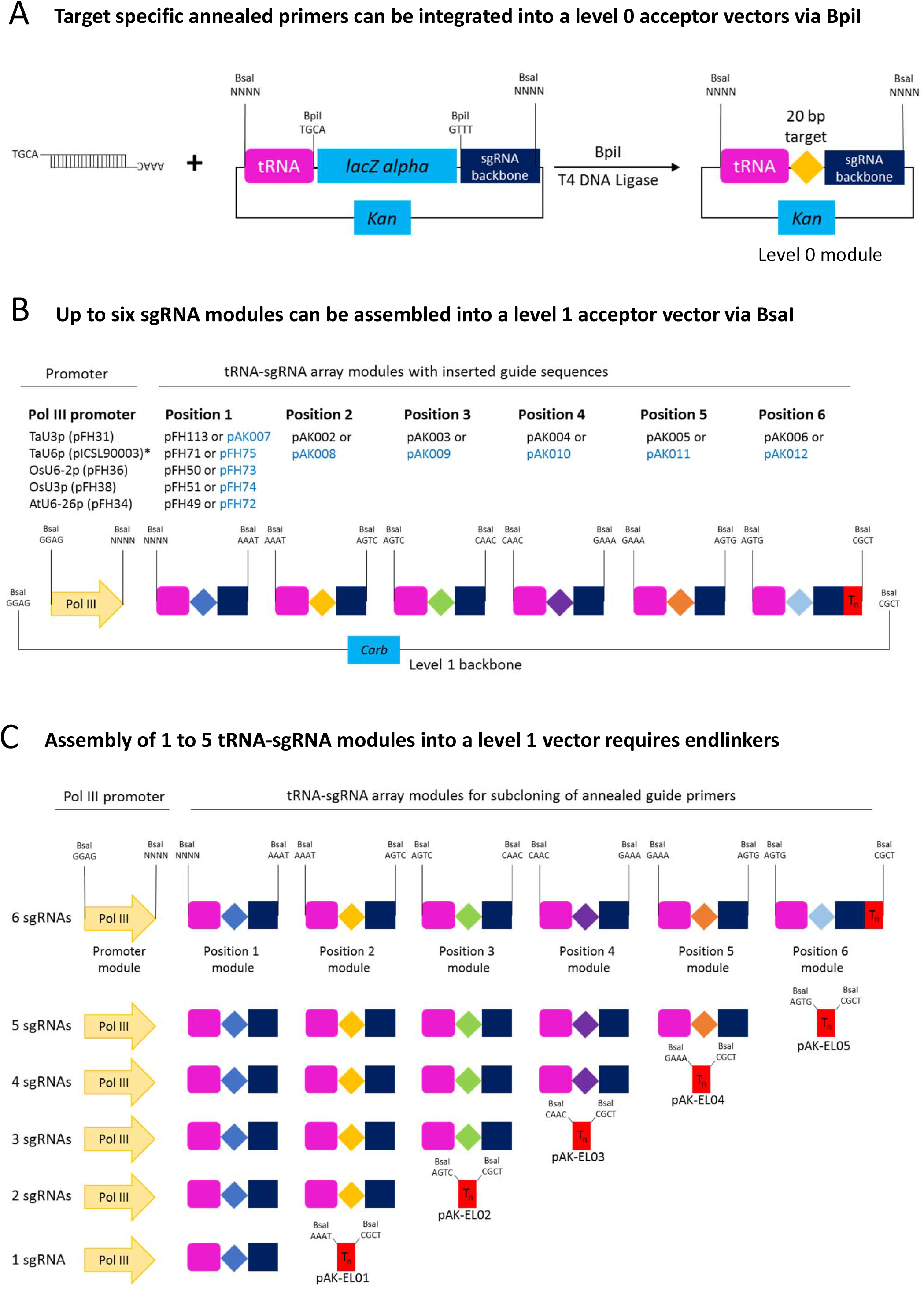
Assembly of level 1 polycistronic tRNA-sgRNA expression modules to be used with SpCas9. During the first step (**a**) annealed complementary oligos encoding the guide are inserted into a level 0 acceptor with the tRNA-sgRNA backbone using BpiI. During the second step (**b**), tRNA-sgRNA modules are fused with the respective Pol III promoter using BsaI. Assembly of less than six tRNA-sgRNA modules into a level 1 vector requires respective endlinkers (**c**). pFH and pAK constructs shown in black font carry the improved sgRNA backbone [39], while the ones shown in blue font – the classic sgRNA backbone [40]. *This is a published module [37].

As stable transformation continues to be a major bottleneck for genome editing applications in many plants, including a major crop like wheat [38], it is advantageous to be able to test CRISPR/Cas constructs for activity in a transient expression system, such as protoplasts, before initiating an often lengthy and labour intensive stable transformation procedure. Using the wheat protoplast system, we have compared activity of three different wheat codon optimised SpCas9 variants (Fig. 6), which mostly differ in their C-terminal NLSs. The fact that one of the constructs (pFH66) performed better than the other two (Fig. 6b) could be due to the histone H2B NLS, located at the C-terminus of the pFH66 variant, being more efficient at importing Cas9 into the nucleus as compared to the SV40 or nucleoplasmin NLS present in the other two constructs. A possible link between different NLS versions and Cas9 activity was previously reported in Arabidopsis [9].

The tRNA-gRNA system for Cas9 multiplexing has proven to work in monocots [6, 35] as well as in dicots [36]. In dicots, it was shown that fusing tRNAs with the optimised sgRNA backbone [39] increased editing efficiencies [36]. In monocots however, only the classic sgRNA backbone [40] was so far used in tRNA-sgRNA arrays. The wheat protoplast assay has allowed us to verify the functionality of the tRNA-sgRNA array, carrying the optimised sgRNA backbone, in a monocot species. Our results suggest that the tRNA-sgRNA array, assembled using the optimised sgRNA backbone, could also result in higher CRISPR/Cas efficiencies in stably transformed monocot plants, in particular, wheat.

**Fig. 6.**
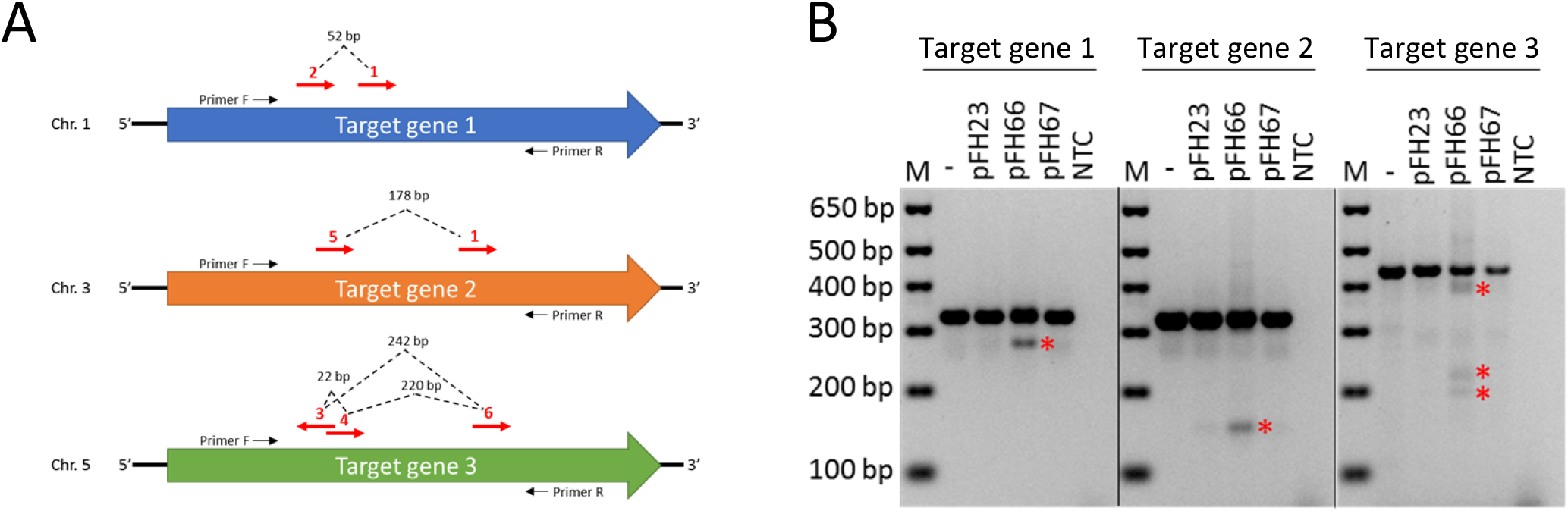
Testing of tRNA-sgRNA CRISPR/Cas constructs in wheat protoplasts. Three wheat genes (**a**) were targeted: Target gene 1 (homoeologues *TraesCS1A02G338200*, *TraesCS1B02G350600* and *TraesCS1D02G340400*), Target gene 2 (homoeologues *TraesCS3A02G289300*, *TraesCS3B02G323900* and *TraesCS3D02G289100*) and Target gene 3 (homoeologues *TraesCS5A02G116500*, *TraesCS5B02G117800* and *TraesCS5D02G129600*). Red arrows illustrate positions of the 20 bp sgRNA target sites. Dashed lines illustrate expected CRISPR/Cas-induced deletions. (**b**) Wheat protoplasts were co-transformed with a level 1 construct carrying one of the three SpCas9 variants (pFH23, pFH66 or pFH67) and the level 1 construct (pFH94) carrying the six sgRNAs (**a**) in a tRNA-sgRNA array. Genotyping by PCR has revealed DNA shifted bands (marked by red asterisks) corresponding to amplicons carrying CRISPR/Cas-induced deletions of expected sizes. ‘M’ is the DNA marker; ‘NTC’ is the no template control.

## Conclusions

We believe the presented modular cloning kit will become a valuable addition to the range of already available GG modules [7, 9, 10] and expect that plant researchers, working with both monocots and dicots, will find the presented molecular tools useful for various genome editing applications. We also believe our study will contribute towards wider adoption of the GG modular cloning system by plant researchers and consequently facilitate exchange of standardised molecular cloning parts across the research community.

## Methods

### DNA construct assembly

All PCR amplifications were performed using Q5® DNA Polymerase (New England Biolabs) according to the manufacturer’s instruction. All GG cut-ligation reactions were performed according to the described protocol (Additional file 1: section 1).

All ligations were transformed into One Shot™ TOP10 chemically competent *E. coli* (Thermo Fisher Scientific) and constructs were verified by sequencing (Eurofins Genomics).

Specific details related to assembly of all GG modules reported in this study are provided (Additional file 1: section 4). Sequences of all PCR primers used in the study are provided (Additional file 1: Table S4). All DNA constructs generated as part of this study were deposited with Addgene (www.addgene.org) with Addgene IDs indicated for each construct (Additional file 2: Table S5). Sequence information of all constructs can be found in GenBank (.gb) files (Additional files 4-105).

### Protoplast assay

Protoplasts were isolated from 10 day old, etiolated wheat seedlings (cv. Cadenza) as previously described [41] with some modifications. Cellulase R10 and Macerozyme R10 were obtained from Duchefa Biochemie (Haarlem, the Netherlands) and the enzymatic digestion was performed at 26 °C for 4 h. Subsequently, 50,000 protoplasts in a volume of 100 μL were transformed with 20 μg of each plasmid (2 μg/μL) purified using the Plasmid Maxi kit (Qiagen, Germany). One batch of protoplasts was treated with an equivalent amount of water and used as the negative (untransformed) control. Then, protoplasts were cultured in 6-well plates for 60 hours. DNA was isolated and CRISPR/Cas-induced mutations were detected by PCR amplification of the target genes using the DreamTaq DNA polymerase (ThermoFisher Scientific) and primer pairs FH432/FH433 (Target gene 1), FH436/FH437 (Target gene 2) and FH440/FH441 (Target gene 3) (Additional file 1: Table S4).

## Supporting information

GenBank DNA construct files

## Abbreviations

CRISPR/Cas: Clustered regularly interspaced short palindromic repeats/CRISPR-associated
crRNA: CRISPR RNA
DSB: Double-strand break
GG: Golden Gate
HDR: Homology-directed repair
MoClo: Modular cloning
PAM: Protospacer adjacent motif
sgRNA: Single guide RNA

## Acknowledgments

We thank Lucy Hyde (University of Bristol) for her contribution of managing the vector database and her help at establishing the protoplast protocol. We thank Robert Hoffie, Ingrid Otto and Nagaveni Budhagatapalli (IPK Gatersleben) for helpful advice on the protoplast assay.

## Authors’ contributions

VN conceived the idea of the manuscript; FH designed the pFH-constructs and performed the experiments; AK designed the pAK-constructs and assisted with cloning; LSL performed protoplast experiments together with FH and assisted with cloning. VN and FH wrote the manuscript.

## Availability of data and materials

The majority of materials are available via Addgene (www.addgene.org) as specified in the manuscript. The rest of materials can be requested by contacting the corresponding author.

## Ethics approval and consent to participate

Not applicable.

## Consent for publication

Not applicable.

