## Supplementary Methods for "A modular cloning toolkit for genome editing in plants"

United Kingdom

**Supplementary methods**

1. **General Golden Gate (GG) cut-ligation reaction** **protocol**
(1) Set up cut-ligations using the following recipe:

| 1 µl | Vector backbone (adjusted to 100 ng/µl) |
| --- | --- |
| 1 µl | Each assembly piece (vectors: adjusted to 100 ng/µl; annealed primers: 10 µM each) |
| 1.5 µl | 10x T4 DNA Ligase buffer (New England Biolabs) |
| 1.5 µl | BSA (1 mg/ml, New England Biolabs) |
| 1 µl  1 µl | BpiI (for assembly of level 0 and level 2 constructs; 10 U/µl, Thermo Fisher Scientific) **or** BsaI-HF®v2 (for assembly of level 1 constructs; 20 U/µl, New England Biolabs)  T4 DNA Ligase (400000 U/µl, New England Biolabs) |
| to 15 µl | ddH_2_O |

(2) Perform GG cut-ligation reaction in a thermocycler using the following program:

| 37 ˚C | 20 s  } 50x |
| --- | --- |
| 37 ˚C | 3 min |
| 16 ˚C | 4 min |
| 50 ˚C | 5 min |
| 80 ˚C | 5 min |
| 16 ˚C | store |

(3) Transform 2 µl of the ligation reaction into One Shot™ TOP10 Chemically Competent *E. coli* (Thermo Fisher Scientific) cells and plate on selection plates.
(4) The following day, pick white colonies and set up overnight cultures.
(5) Extract plasmid DNA using the Qiagen miniprep kit and verify constructs by restriction digestion and Sanger sequencing.
2. **Assembly of CRISPR/Cas constructs with sgRNA/crRNA expressed under individual Pol III promoters**

This protocol will provide a detailed strategy for cloning of a L2 vector containing a selectable marker cassette, a transcriptional unit for a nuclease and transcriptional unit(s) for the corresponding sgRNA/crRNA (Fig. 1). For a general overview how GG cloning works, please refer to Weber et al. (2011). A timeline for assembly of a generic level 2 CRISPR/Cas construct can be found in the Figure S1.

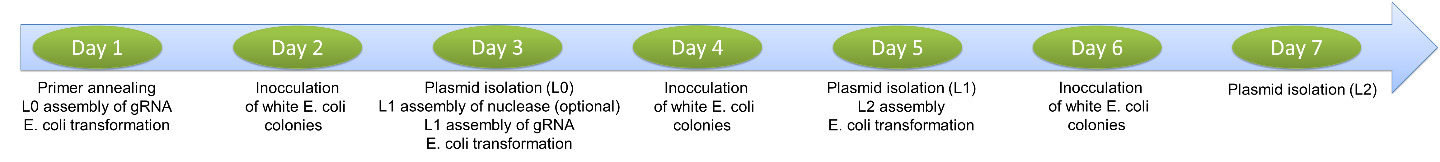

Figure S1 A timeline for assembly of a level 2 CRISPR/Cas construct.

Step 1: Subcloning of your target into the correct acceptor vector

(1) Choose the correct subcloning vector from Table S1 for your target sequence depending on the Pol III promoter that you intend to use in your experiment (wheat *TaU3* promoter for monocots, Arabidopsis *AtU6-26* promoter for dicots) and depending which nuclease you want to use.

Table S1

A list of modules encoding guide RNA backbones with matching CRISPR/Cas nucleases and Pol III promoters

| **Vector ID** | **Guide RNA backbone** | **Works with these nucleases** | **Works with this promoter module** |
| --- | --- | --- | --- |
| pFH85 | sgRNA classic backbone | SpCas9, SpCas9-NG, ScCas9, nScCas9, xCas9, all base editors | Wheat *TaU3p* (pFH33) |
| pFH86 | sgRNA improved backbone [1] | SpCas9, SpCas9-NG, ScCas9, nScCas9, xCas9, all base editors | Wheat *TaU3p* (pFH33) |
| pFH87 | StCas9 sgRNA backbone | StCas9 | Wheat *TaU3p* (pFH33) |
| pFH88 | SaCas9 sgRNA backbone | SaCas9 | Wheat *TaU3p* (pFH33) |
| pFH89 | LbCas12a crRNA backbone | LbCas12a | Wheat *TaU3p* (pFH33) |
| pFH90 | FnCas12a crRNA backbone | FnCas12a, all Cms1 nucleases | Wheat *TaU3p* (pFH33) |
| pFH99 | sgRNA classic backbone | SpCas9, SpCas9-NG, ScCas9, xCas9, all base editors | Arabidopsis *AtU6-26p* (pFH35) |
| pFH100 | sgRNA improved backbone [1] | SpCas9, SpCas9-NG, ScCas9, xCas9, all base editors | Arabidopsis *AtU6-26p* (pFH35) |
| pFH101 | StCas9 sgRNA backbone | StCas9 | Arabidopsis *AtU6-26p* (pFH35) |
| pFH102 | SaCas9 sgRNA backbone | SaCas9 | Arabidopsis *AtU6-26p* (pFH35) |
| pFH103 | LbCas12a crRNA backbone | LbCas12a | Arabidopsis *AtU6-26p* (pFH35) |
| pFH104 | FnCas12a crRNA backbone | FnCas12a, all Cms1 nucleases | Arabidopsis *AtU6-26p* (pFH35) |

(2) Find suitable target sequences in your gene of interest.

*Note: The only requirement for your target sequence is that it lacks BpiI (GAAGAC) and BsaI (GGTCTC) recognition sites as this will interfere with GG cloning*

(3) Order two complementary primers encoding your target sequence and the correct overhangs depending on the subcloning vector that you will use (Table S2).

Table S2

Primer design for inserting guide sequences into respective guide RNA backbone modules

| **Vector ID** | **Guide RNA backbone** | **FW primer** | **Rev primer** |
| --- | --- | --- | --- |
| pFH85 | sgRNA classic backbone | AGCA (N)_20_ | AAAC (N)_20 reverse complement_ |
| pFH86 | sgRNA improved backbone [1] | AGCA (N)_20_ | AAAC (N)_20 reverse complement_ |
| pFH87 | StCas9 sgRNA backbone | AGCA (N)_20_ | AAAC (N)_20 reverse complement_ |
| pFH88 | SaCas9 sgRNA backbone | AGCA (N)_20_ | AAAC (N)_20 reverse complement_ |
| pFH89 | LbCas12a crRNA backbone | AGAT (N)_23_ | AAAA (N)_23 reverse complement_ |
| pFH90 | FnCas12a crRNA backbone | AGAT (N)_23_ | AAAA (N)_23 reverse complement_ |
| pFH99 | sgRNA classic backbone | ATTG (N)_20_ | AAAC (N)_20 reverse complement_ |
| pFH100 | sgRNA improved backbone [1] | ATTG (N)_20_ | AAAC (N)_20 reverse complement_ |
| pFH101 | StCas9 sgRNA backbone | ATTG (N)_20_ | AAAC (N)_20 reverse complement_ |
| pFH102 | SaCas9 sgRNA backbone | ATTG (N)_20_ | AAAC (N)_20 reverse complement_ |
| pFH103 | LbCas12a crRNA backbone | AGAT (N)_23_ | AAAA (N)_23 reverse complement_ |
| pFH104 | FnCas12a crRNA backbone | AGAT (N)_23_ | AAAA (N)_23 reverse complement_ |

*Note: The corresponding promoter modules contain already the correct transcriptional start nucleotide for expression of the guides (A in case of U3 promoters, G in case of U6 promoters) at their 3’-end to allow transcription of all guides independent of their first nucleotide. If your target sequence starts with the correct transcriptional start nucleotide already, consider to not add it to your primer sequence (example: for pFH85, you would just order two primers AGCA (N)_2-20_ and AAAC (N)_20-2 reverse complement_). This does not apply to the Cas12a/Cms1 crRNAs, as the target sequence is anyway at the 3’end of the crRNA sequence.*

(4) Adjust primer concentrations to 10 µM and mix the forward and reverse in a 1:1 ratio, incubate for 5 min at room temperature
(5) Integrate your annealed primers into the chosen subcloning vector using a GG cut-ligation reaction via BpiI (Thermo Fisher Scientific) (Figs. 2a and 3a).
(6) Transform 2 µL of the GG reaction into competent cells and plate the cells on selection plates containing kanamycin, IPTG and X-Gal.
(7) Inoculate a white colony, prepare plasmid DNA and verify correct integration of your annealed primer by sequencing with primer FH32 (Table S4).

Step 2: Generation of transcription unit for guide RNA expression

(8) Combine the subcloned target module with the correct Pol III promoter module (see Table S1) into a L1 backbone (see Additional file 3: Table S6) using a GG cut-ligation reaction via BsaI-HF®v2 (New England Biolabs) (Figs. 2b and 3b).

*Note: In most cases, the most suitable L1 backbone is pICH47751* [2] *(Addgene ID: 48002), which will place your promoter-gRNA unit in Position 3 in the final L2 vector. If you want to add several separate promoter-gRNA units in the final L2 constructs, then you have to clone them separately in different L1 backbones in this step (e.g. Position 3, 4, 5).*

(9) Transform 2 µL of the GG reaction into competent cells and plate the cells on selection plates containing carbenicillin, IPTG and X-Gal.
(10) Inoculate a white colony, prepare plasmid DNA and verify correct assembly by sequencing with primers Lvl1_F(0229) and Lvl1_R(0230) (Table S4).

Step 3: Generation of a multi-expression unit L2 construct (Fig. 1)

(11) Combine your L1 sgRNA/crRNA construct(s) with a L1 resistance marker transcription unit (Tables S5 and S6), a suitable nuclease L1 module (Additional file 2: Table S5) and the correct endlinker module (Additional file 3: Table S6; in case of one promoter-gRNA unit: pICH41766 ([2] (Addgene ID #48018)) into a suitable L2 backbone (Additional file 3: Table S6) using a GG cut-ligation reaction via BpiI (Thermo Fisher Scientific).
(12) Transform 2 µL of the GG reaction into competent cells and plate the cells on selection plates.
(13) Inoculate a white colony, prepare plasmid DNA and verify correct assembly by suitable restriction digest and sequencing.
3. **CRISPR/Cas construct assembly for multiple targets using the polycistronic tRNA-sgRNA system**

This protocol will provide a detailed strategy for cloning of a L2 vector containing a resistance cassette, a transcriptional unit for a nuclease and a set of up to six sgRNAs using the polycistronic sgRNA-tRNA multiplex system [3] (Fig. 4). For a general overview how the GG cloning works, please refer to Weber et al. [2].

Step 1: Subcloning of your targets into the correct acceptor vector

(1) Choose the correct subcloning vector for sgRNA that you intend to clone at the 5’-end of the polycistronic tRNA-sgRNA (position 1) depending on the Pol III promoter that you intend to use in your experiment and depending which sgRNA backbone you prefer according to Table S3 and Fig. 5b.

Table S3

Pol III promoter modules and matching position 1 modules in the tRNA-sgRNA array

| Pol III promoter module | | Matching position 1 module in the  tRNA-sgRNA array | |
| --- | --- | --- | --- |
| Vector ID | Description | SpCas9 sgRNA improved backbone [1] | SpCas9 sgRNA classic backbone [4] |
| pFH31 | Wheat *TaU3p* | pFH113 | pAK007 |
| pICSL90003  [5] | Wheat *TaU6p* | pFH71 | pFH75 |
| pFH36 | Rice *OsU6-2p* | pFH50 | pFH73 |
| pFH38 | Rice *OsU3p* | pFH51 | pFH74 |
| pFH34 | Arabidopsis *AtU6-26p* | pFH49 | pFH72 |

(2) Find suitable target sequences in your gene of interest.

*Note: The only requirement for your target sequence is that it lacks BpiI (GAAGAC) and BsaI (GGTCTC) recognition sites as this will interfere with GG cloning*

(3) Order two complementary primers for each of your target sequences as following:

5’ TGCA (N)_20_ 3’

5’ AAAC (N)_20 reverse complement_ 3’.

(4) Adjust primer concentrations to 10 µM and mix the forward and reverse in a 1:1 ratio, incubate for 5 min at room temperature
(5) Integrate your annealed primers into the different subcloning vectors according to the position in which you want your target using a GG cut-ligation reaction via BpiI (Thermo Fisher Scientific) (Fig. 5a).
(6) Transform 2 µL of the GG reaction into competent cells and plate the cells on selection plates containing kanamycin, IPTG and X-Gal.
(7) Inoculate a white colony, prepare plasmid DNA and verify correct integration of your annealed primer by sequencing with primer FH32 (Table S4).

Step 2: Generation of a polycistronic tRNA-sgRNA transcription unit

(8) Combine the subcloned target module with the correct promoter module (see Table S3 and Fig. 5b) and a suitable endlinker module (only necessary, if less than six gRNAs are cloned; Fig. 5c) into a L1 backbone (Additional file 3: Table S6) using a GG cut-ligation reaction via BsaI-HF®v2 (New England Biolabs) (Fig. 5b,c).

*Note: In most cases, the most suitable L1 backbone is pICH47751* [2] (*Addgene ID: 48002*)*, which will place your promoter-gRNA unit in Position 3 in the final L2 vector. If you want to add several separate polycistronic tRNA-sgRNA units in the final L2 constructs, then you have to clone them separately in different L1 backbones in this step (e.g. Position 3, 4, 5).*

(9) Transform 2 µL of the GG reaction into competent cells and plate the cells on selection plates containing carbenicillin, IPTG and X-Gal.
(10) Inoculate a white colony, prepare plasmid DNA and verify correct assembly by sequencing with primers Lvl1_F(0229) and Lvl1_R(0230).

Step 3: Generation of a multi-expression unit L2 construct (Fig. 4)

(11) Combine your L1 polycistronic tRNA-sgRNA construct with a L1 resistance marker transcription unit (Tables S5 and S6), a suitable SpCas9-based L1 module (Tables S5 and S6) and the correct endlinker module (in case of one polycistronic tRNA-sgRNA unit: pICH41766 [2] Addgene ID #48018) into a suitable L2 backbone (Additional file 3: Table S6) using a GG cut-ligation reaction via BpiI (Thermo Fisher Scientific).
(12) Transform 2 µL of the GG reaction into competent cells and plate the cells on selection plates.
(13) Inoculate a white colony, prepare plasmid DNA and verify correct assembly by suitable restriction digest and sequencing.

Table S4

Primers used in this study

| **Primer name** | **Sequence (5’-3’)** |
| --- | --- |
| FH1 | ttgaagacaaAATGGCACCGAAGAAGAAGC |
| FH2 | ttgaagacaaAAGCTTAACTTTTCTTCTTCTTGTCGG |
| FH3 | ttgaagacaaAATGGGGCCGAAGAAGAAGAGG |
| FH4 | ttgaagacaaAAGCTTAGACTTTGCGCTTCTTCTTGGGA |
| FH5 | ttgaagacaaAATGAAGAGGAACTACATCCTCG |
| FH6 | ttgaagacaaAAGCTCAAACCTTCCTCTTCTTCTTAG |
| FH7 | ttgaagacaaAATGTCTGATCTCGTGCTCGG |
| FH8 | ttgaagacaaAAGCTCAAACCTTCCTCTTCTTCTTAGG |
| FH9 | ttgaagacatctcaGCTTTCCAGGCCTCCCAGCTTT |
| FH10 | ttgaagacaactcgAGCGGGGAGCTCAAGCCTATACTGTACTT |
| FH11 | ttgaagacatctcaGCTTGTTCGAGTATTATGGCATTGG |
| FH12 | ttgaagacaactcgAGCGACAACTACAAGTGTTTTACTCCTCA |
| FH13 | ttgaagacaaGGAGCACGTCAGTGTTTGGTTTCCAC |
| FH14 | ttgaagacaaCATtGCAGGATCTGCAGAAGATTAGC |
| FH15 | ttgaagacaaGGAGAAAAATTACGGATATGAATATAGGC |
| FH16 | ttgaagacaaCATtGCTGCACATACATAACATATCAAG |
| FH17 | ttgaagacaaAATGAGCATCTACCAGGAGTTCGTC |
| FH18 | ttgaagacaaAAGCTCACTTTTTCTTTTTTGCCTGGCCGGCC |
| FH19 | ttgaagacaaAATGTCCGAAGTCGAGTTTTCC |
| FH20 | ttgaagacaaAAGCTTAGACTTTCCTCTTCTTCTTGGG |
| FH21 | ttgaagacaactcaGGAGAAGCTTCGTTGAACAACGGA |
| FH22 | ttgaagacaactcgAATCACTACTTCGACTCTAGCTGT |
| FH23 | ttgaagacaactcaGGAGCATGAATCCAAACCACACGGAG |
| FH24 | ttgaagacaactcgGCTTCTTGGTGCCGCGC |
| FH25 | ttgaagacaactcaGGAGGGATCATGAACCAACGGCC |
| FH26 | ttgaagacaactcgAACACAAGCGACAGCGCG |
| FH27 | ttgaagacaactcaGGAGAAGGAATCTTTAAACATACGAACAG |
| FH28 | ttgaagacaactcgGCCACGGATCATCTGCA |
| FH29 | ttgaagacaaGTACTGGTGCTACCAGCAAATG |
| FH30 | ttgaagacaaGTACAAGACGCCTGTCTTGTGCTAT |
| FH32 | GCAATGTAACATCAGAGATTTTGA |
| FH33 | ttgaagacaaAATGTCCATCTACCAAGAGTTCGT |
| FH34 | ttgaagacaaTTTGGCGATCAGCTCTTGCT |
| FH35 | ttgaagacaaCAAAAAGACCGAGAAGGCGAAGTAC |
| FH36 | ttgaagacaaGGTTTTCATGCGGTCGTTGCC |
| FH37 | ttgaagacaaAACCAACTACCACGATAAGCTGG |
| FH38 | ttgaagacaaAAGCTCAGACCTTCCTCTTCTTCTTTG |
| FH39 | ttgaagacaaAATGGACCACTACCTCGACATC |
| FH40 | ttgaagacaaAAGCTTAGAACCACGGCACGAAGC |
| FH60 | cacattgaagacaaAATGGCTCCTAAGAAGAAGCGG |
| FH61 | cacattgaagacaaATCTTCCGCGCGCTTTTC |
| FH62 | ttgaagacaaAGATTACAAAGGGGTGAAGAAGTTGTT |
| FH63 | ttgaagacaaTAACAGGTTGACAAAACCAGTTG |
| FH64 | ttgaagacaaGTTAAAGACGAAATACACGTCCATTG |
| FH65 | ttgaagacaaCTCGGCCTTTTTGAACTGG |
| FH66 | ttgaagacaaCGAGGACGAAAAATTGGACAAGGTC |
| FH67 | ttgaagacaaAAGCTCACTTCTTTTTCTTAGCCTGTC |
| FH68 | AATGGCCCCAAAGAAGAAACGGAAGGTTATGGAGAAGTACAAGAT |
| FH69 | GGTTATCTTGTACTTCTCCATAACCTTCCGTTTCTTCTTTGGGGC |
| FH70 | ttgaagacaaAACCAAGACGATCAGATTTAAGCTGCTG |
| FH71 | ttgaagacaaCTCAAACGCAAATGTGATCTCGT |
| FH72 | ttgaagacaaTGAGGACACAAAGAAGAAGGGCACG |
| FH73 | ttgaagacaaTCAGACCTTCGAACCCTTTCAGG |
| FH74 | CTGAACGACCCAGATAAGGTCGCTGCGTTCAACATTGCGAAGCGCGGTTTCGAGGATCTTCAGAAGTACAAGTAA |
| FH75 | AAGCTTACTTGTACTTCTGAAGATCCTCGAAACCGCGCTTCGCAATGTTGAACGCAGCGACCTTATCTGGGTCGT |
| FH137 | ttgaagacaaAATGGCACCTAAAAAAAAGAGG |
| FH138 | ttgaagacaaCTCGAAGAGCGTGAGTGTGAG |
| FH139 | ttgaagacaaCGAGGACCGGGAAATGATTGAAGA |
| FH140 | ttgaagacaaTATCTTCAGGGGAACCCTTGA |
| FH141 | ttgaagacaaGATAATGAGCAAAAACAGCTCTTC |
| FH142 | ttgaagacaaAAGCTCATTTCTTCTTTTTCGCTTG |
| FH158 | ttgaagacaaGGTTTTCATCTTTCTGTAAGCCTCGA |
| FH159 | ttgaagacaaAACCCTCGAAACCTTGGATATC |
| FH239 | ttgaagacaaAATGGCCCCAAAGAAGAAG |
| FH240 | ttgaagacaaAAGCTTACTTTTTCTTTTTTGCCTGG |
| FH275 | ttgaagacaactcgCAATCACTACTTCGACTCTAGCTG |
| FH276 | ttgaagacaactcgTGCTTCTTGGTGCCGCGC |
| FH277 | ttgaagacaactcgCAACACAAGCGACAGCGCG |
| FH278 | ttgaagacaactcgTGCCACGGATCATCTGC |
| FH285 | ttgaagacaaGGAGTGGCAGACATACTGTCCCACA |
| FH286 | ttgaagacaaCATtAAGCTTAGCTCTTACCTGTTTTCG |
| FH287 | ttgaagacaaAATGAGCAAGCTGGAGAAGTTTACA |
| FH303 | GATGGTCTCAgattGAACAAAGCACCAGTGGTC |
| FH304 | GACAATGACGTCCGGGAG |
| FH305 | GATGGTCTCAtgttGAACAAAGCACCAGTGG |
| FH306 | GATGGTCTCAtggcAACAAAGCACCAGTGGTC |
| FH307 | ttgaagacaaGGAGCTAGCATACTCGAGGTCATTCAT |
| FH308 | ttgaagacaaCATtGGCTTCTACCTACAAAAAAGCTC |
| FH309 | ttgaagacaaGGAGGGGGCCCAATATAACAACGAC |
| FH310 | ttgaagacaaCATtTTGTCGAGTCAACAATCACAGAT |
| FH311 | ttgaagacaaAATGGATAAGAAGTACTCTATCGGAC |
| FH312 | ttgaagacaaAAGCTCAGGGCCCTACTTTACGT |
| FH315 | ttgaagacaaCGACTTCGGAAGAACCACC |
| FH316 | ttgaagacaaGTCGAGTTTTCCCATGAGTACTG |
| FH323 | GATGGTCTCActtgAACAAAGCACCAGTGGTCTAG |
| FH348 | ATTGGGCTGGcCATTGGCACC |
| FH349 | AGAGTATTTCTTCTCCATTTGAGACC |
| FH352 | ttgaagacaaAAGCCTACACCTTGCGCTTCTTCTTT |
| FH360 | tgcaTGGGGCATTGGATTGGGATT |
| FH361 | aaacAATCCCAATCCAATGCCCCA |
| FH362 | tgcaCTTTATCCAGGACCTGGAGA |
| FH363 | aaacTCTCCAGGTCCTGGATAAAG |
| FH364 | tgcaAACGTCCTCTGCATCCGCCT |
| FH365 | aaacAGGCGGATGCAGAGGACGTT |
| FH366 | tgcaGCAGAGGACGTTGACGACGC |
| FH367 | aaacGCGTCGTCAACGTCCTCTGC |
| FH368 | tgcaGAATGTAGAGGCAACTGCAG |
| FH369 | aaacCTGCAGTTGCCTCTACATTC |
| FH370 | tgcaGACATGGAGGTCCTCTACGT |
| FH371 | aaacACGTAGAGGACCTCCATGTC |
| FH406 | ttgaagacaaAATGGACAAGAAGTACTCCAT |
| FH407 | ttgaagacaaAAGCCTACACCTTCCTCTTCTTCTTGG |
| FH411 | CTGGGTCTCAattgTTGTCTTCGCAGCTGGC |
| FH412 | GTCAGGCGGAATGGCACT |
| FH413 | CTGGGTCTCAattgTAATTTCTACTAAGTGTAGATTTGTCTTCGC |
| FH414 | GTCAGGCGGAATGGCACT |
| FH415 | CTGGGTCTCAattgTAATTTCTACTGTTGTAGATTTGTCTTCGC |
| FH418 | ttgaagacaaCAATAGAATACTTTTTCTCAGAACCACCA |
| FH419 | ttgaagacaaATTGGGCTGGCCATTGGC |
| FH432 | AGCTGAGGCACGAAAAAC |
| FH433 | TCTTATCCGAGCATAGGGTAT |
| FH436 | CAGAGTTCATTGGGTGCTCAC |
| FH437 | CCTCTCGTAAGCCATAACAAT |
| FH440 | CCAATGGGAATCTCTCGGC |
| FH441 | GTAGAAGCGGTCGGAGAGGTC |
| FH443 | ttGGTCTCaggagTGACCCGGTCGTGCCCCT |
| FH444 | ttGGTCTCaagcgCAGTAACATAGATGACACCGCGCG |
| Lvl1_F(0229) | GAACCCTGTGGTTGGCATGCACATAC |
| Lvl1_R(0230) | CTGGTGGCAGGATATATTGTGGTG |

4. **Assembly of cloning toolkit components generated in this study**

All vector sequences can be found as GenBank supplementary files (Additional files 4-105). The following people kindly provided vectors which we used for cloning our toolkit components: Sylvestre Marillonnet (IPB, Germany), Holger Puchta (KIT, Germany), Daniel Voytas (U. of Minnesota, USA), Jonathan Jones (TSL, UK), Mark Youles (TSL, UK), Nicola Patron (EI, UK), Eduard Akhunov (KSU, USA), Seeichi Toki (Yokohama City U., Japan), Yiping Qi (U. of Maryland, USA), Matthew Begemann (Benson Hill, USA), Keith Edwards (U. of Bristol, UK), Alison Huttly (Rothamsted Research, UK), Damiano Martignago (Rothamsted Research, UK), David Liu (MIT, USA), Andreas Weber (U. of Dusseldorf, Germany), Feng Zhang (Broad Institute, USA), Akihiko Kondo (Kobe U., Japan), Joseph Jacobson (MIT, USA), Caixia Gao (CAS, China) and Caroline Sparks (Rothamsted Research, UK).

**Level 0 CRISPR/Cas nuclease CDS constructs**

pFH13: The wheat codon optimised *SpCas9* gene was amplified in fragments from the vector pA9Cas9 [6] using primers FH137/FH138, FH139/FH140 and FH141/FH142. The fragments were subcloned into pCR-BluntII-Topo using the Zero Blunt™ TOPO™ PCR Cloning Kit (ThermoFisher Scientific) and then assembled into pICH41308 [2] (Addgene ID #47998) using a GG cut-ligation reaction via BpiI.

pFH14: The Arabidopsis codon optimised *SaCas9* gene was amplified from the vector pDE-Sa-Cas9 [7] using primers FH5/FH6 and the fragments were subcloned into pCR-BluntII-Topo. The *SaCas9* gene was then assembled into pICH41308 [2] (Addgene ID #47998) using a GG cut-ligation reaction via BpiI.

pFH15: The Arabidopsis codon optimised *StCas9* gene was amplified in fragments from the vector pDE-St-Cas9 [7] using primers FH7/FH158 and FH8/FH159 and the fragments were subcloned into pCR-BluntII-Topo. The *StCas9* gene was then assembled into pICH41308 [2] Addgene ID #47998) using a GG cut-ligation reaction via BpiI.

pFH16: The rice codon optimised *FnCas12a* gene was amplified in fragments from the vector pPZP-FnCas12a-OsOpt [8] using primers FH33/FH34, FH35/FH36 and FH37/FH38. The fragments were subcloned into pCR-BluntII-Topo and the *FnCas12a* gene was then assembled into pICH41308 [2] (Addgene ID #47998) using a GG cut-ligation reaction via BpiI.

pFH17: The rice codon optimised *LbCas12a* gene was amplified in fragments from the vector pYPQ230 [9] (Addgene ID #86210) using primer pairs FH60/FH61, FH62/FH63, FH64/FH65 and FH66/FH67. The fragments were subcloned into pCR-BluntII-Topo using the Zero Blunt™ TOPO™ PCR Cloning Kit (ThermoFisher Scientific) and then assembled into pICH41308 [2] (Addgene ID #47998) using a GG cut-ligation reaction via BpiI.

pFH18: The monocot optimised *SmCms1* gene was amplified in fragments from the vector SmCms1 (kindly provided by Benson Hill, USA) [10] using primers FH70/FH71 and FH72/FH73 and the fragments were subcloned into pCR-BluntII-Topo. The *SmCms1* gene was then assembled into pICH41308 [2] (Addgene ID #47998) using a GG cut-ligation reaction via BpiI using the subcloned fragments and the annealed primer pairs FH68/FH69 and FH74/FH75. Benson Hill (USA) is the source of the *SmCms1* vector/sequence and the license to the licensed patent rights.

pFH19: The monocot optimised *MiCms1* gene was synthesised in two parts with BpiI overhangs by Twist Bioscience (San Francisco, USA) based on the published sequence [10]. The two parts were assembled into pICH41308 [2] (Addgene ID #47998) using a GG cut-ligation reaction via BpiI. Benson Hill (USA) is the source of the *MiCms1* sequence and the license to the licensed patent rights.

pFH20: The monocot optimised *ObCms1* gene was synthesised in two parts with BpiI overhangs by Twist Bioscience (San Francisco, USA) based on the published sequence [10]. The two parts were assembled into pICH41308 [2] (Addgene ID #47998) using a GG cut-ligation reaction via BpiI. Benson Hill (USA) is the source of the *ObCms1* sequence and the license to the licensed patent rights.

pFH21: The monocot optimised *SuCms1* gene was synthesised in two parts with BpiI overhangs by Twist Bioscience (San Francisco, USA) based on the published sequence [10]. The two parts were assembled into pICH41308 [2] (Addgene ID #47998) using a GG cut-ligation reaction via BpiI. Benson Hill (USA) is the source of the *SuCms1* vector/sequence and the license to the licensed patent rights.

pFH22: The *xCas9-3.7* gene was amplified from the vector pEPOR0SP0011 [11] using primers FH407/FH407. The gel-purified PCR product was assembled into pICH41308 [2] (Addgene ID #47998) using a GG cut-ligation reaction via BpiI.

pFH24: The wheat codon optimised *SpCas9* gene was amplified from the vector pRRes226RRM.486 (kindly provided by Alison Huttly/Damiano Martignago, Rothamsted Research) using primers FH1/FH2. The gel-purified PCR product was assembled into pICH41308 [2] (Addgene ID #47998) using a GG cut-ligation reaction via BpiI.

pFH25: The wheat codon optimised *SpCas9* gene was amplified from the vector pENTR4:Wheat_live_Cas9_P2A_Histone_GFP (kindly provided by Keith Edwards, University of Bristol) using primers FH3/FH4. The gel-purified PCR product was assembled into pICH41308 [2] (Addgene ID #47998) using a GG cut-ligation reaction via BpiI.

pFH32: The human codon optimised *SpCas9-NG* gene was amplified from the vector pX330-SpCas9-NG [12] (Addgene ID #117919) using primers FH239/FH240. The gel-purified PCR product was assembled into pICH41308 [2] (Addgene ID #47998) using a GG cut-ligation reaction via BpiI.

pFH45: The human codon optimised *ABE7.10* A-G base editor gene was amplified in fragments from the vector pCMV-ABE7.10 [13] (Addgene ID #102919) using primer pairs FH19/FH315 and FH20/FH316. The gel-purified PCR products were assembled into pICH41308 [2] (Addgene ID #47998) using a GG cut-ligation reaction via BpiI.

pFH46: The human codon optimised *FnCas12a* gene was amplified from the vector pY004 [14] (Addgene ID #69976) using primers FH17/FH18. The gel-purified PCR product was assembled into pICH41308 [2] (Addgene ID #47998) using a GG cut-ligation reaction via BpiI.

pFH47: The human codon optimised *LbCas12a* gene was amplified from the vector pY016 [14] (Addgene ID #69988) using primers FH287/FH18. The gel-purified PCR product was assembled into pICH41308 [2] (Addgene ID #47998) using a GG cut-ligation reaction via BpiI.

pFH48: The *Csy4* gene was amplified from the vector pDirect_26H [15] (Addgene ID #91150) using primers FH39/FH40. The gel-purified PCR product was assembled into pICH41308 [2] (Addgene ID #47998) using a GG cut-ligation reaction via BpiI.

pFH55: The Arabidopsis codon optimised C-T base editor *nSpCas9-PmCDA* gene was amplified from the vector pDicAID_nCas9-PmCDA_NptII_Della [16] (Addgene ID #91694) using primers FH311/FH312. The gel-purified PCR product was assembled into pICH41308 [2] (Addgene ID #47998) using a GG cut-ligation reaction via BpiI.

pFH76: The wheat codon optimised *ScCas9* gene was synthesised in two parts with BpiI overhangs by Twist Bioscience (San Francisco, USA) based on the published sequence [17]. The two parts were assembled into pICH41308 [2] (Addgene ID #47998) using a GG cut-ligation reaction via BpiI.

pFH77: The wheat codon optimised *enCas9-PolI3M-TBD* gene was synthesised in three parts with BpiI overhangs by Twist Bioscience (San Francisco, USA) based on the published sequence [18]. The three parts were assembled into pICH41308 [2] (Addgene ID #47998) using a GG cut-ligation reaction via BpiI.

pFH79: The wheat codon optimised *A3A-PBE* C-T base editor gene was synthesised in two parts with BpiI overhangs by Twist Bioscience (San Francisco, USA) based on the published sequence [19]. The two parts were assembled into pICH41308 [2] (Addgene ID #47998) using a GG cut-ligation reaction via BpiI.

pFH91: The *nScCas9* construct was created by introduction of the D10A mutation into pFH76 via the Q5® Site-directed Mutagenesis Kit (New England Biolabs) using primers FH348/FH349 according to the supplier’s manual.

pFH92: The nScCas9-ABE7.10 A-G base editor construct was assembled by amplifying the ABE7.10 base editor domain from pFH45 using primers FH315/FH19 and FH418/FH316 and amplifying the nScCas9 from pFH91 using primers FH419/FH352. The first PCR product was subcloned into pCR-BluntII-Topo and assembled with the other two gel-purified PCR products into pICH41308 [2] (Addgene ID #47998) using a GG cut-ligation reaction via BpiI.

**Level 0 Pol II promoter + 5’UTR constructs**

pFH40: The switchgrass *UBIQUITIN1* promoter was amplified from the vector pMOD_B2312 [15] (Addgene ID #91070) using primers FH13/FH14. The gel-purified PCR product was assembled into pICH41295 [2] (Addgene ID #47997) using a GG cut-ligation reaction via BpiI.

pFH41: The parsley *UBIQUITIN4-2* promoter was amplified from the vector pDE-Cas9 [20] (Addgene ID #61433) using primers FH15/FH16. The gel-purified PCR product was assembled into pICH41295 [2] (Addgene ID #47997) using a GG cut-ligation reaction via BpiI.

pFH42: The Cestrum Yellow Leaf Curling Virus *CmYLCV* promoter was amplified from the vector pMOD_C3003 [15] (Addgene ID #91095) using primers FH385/FH286. The gel-purified PCR product was assembled into pICH41295 [2] (Addgene ID #47997) using a GG cut-ligation reaction via BpiI.

pFH56: The rice *Actin1* promoter was amplified from the vector pICSL11100 [21] using primers FH307/FH308. The gel-purified PCR product was assembled into pICH41295 [2] (Addgene ID #47997) using a GG cut-ligation reaction via BpiI.

pFH57: The soybean *UBIQUITIN* promoter was amplified from the vector pMOD_B2121 [15] (Addgene ID #91065) using primers FH309/FH310. The gel-purified PCR product was assembled into pICH41295 [2] (Addgene ID #47997) using a GG cut-ligation reaction via BpiI.

**Level 0 3’UTR + terminator constructs**

pFH43: The pea *RbSC* terminator was amplified from the vector pMOD_C3003 [15] (Addgene ID #91095) using primers FH11/FH12. The gel-purified PCR product was assembled into pAGM9121 [2] (Addgene ID #51833) using a GG cut-ligation reaction via BpiI.

pFH44: The pea *3A* terminator was amplified from the vector pDE-Cas9 [20] (Addgene ID #61433) using primers FH9/FH10. The gel-purified PCR product was assembled into pAGM9121 [2] (Addgene ID #51833) using a GG cut-ligation reaction via BpiI.

**Level 0 Pol III promoter constructs**

pFH31: The wheat *U3* promoter was amplified from the vector pBUN421-GLM [6] using primers FH23/FH24 and assembled into pAGM9121 [2] (Addgene ID #51833) using a GG cut-ligation reaction via BpiI.

pFH33: The wheat *U3* promoter was amplified from the vector pBUN421-GLM [6] using primers FH23/FH276 and assembled into pAGM9121 [2] (Addgene ID #51833) using a GG cut-ligation reaction via BpiI.

pFH34: The Arabidopsis *U6-26* promoter was amplified from the vector pFH6_new [22] (Addgene ID #105866) using primers FH21/FH22 and assembled into pAGM9121 [2] (Addgene ID #51833) using a GG cut-ligation reaction via BpiI.

pFH35: The Arabidopsis *U6-26* promoter was amplified from the vector pFH6_new [23] (Addgene ID #105866) using primers FH21/FH275 and assembled into pAGM9121 [2] (Addgene ID #51833) using a GG cut-ligation reaction via BpiI.

pFH36: The rice *U6-2* promoter was amplified from the vector pUC19_OsU6_SpCas9_sgRNA [8] using primers FH25/FH26 and assembled into pAGM9121 [2] (Addgene ID #51833) using a GG cut-ligation reaction via BpiI.

pFH37: The rice *U6-2* promoter was amplified from the vector pUC19_OsU6_SpCas9_sgRNA [8] using primers FH25/FH277 and assembled into pAGM9121 [2] (Addgene ID #51833) using a GG cut-ligation reaction via BpiI.

pFH38: The rice *U3* promoter was amplified in fragments from the vector pPZP-FnCas12a-OsOpt [8] using primers FH27/FH30 and FH28/FH29. Both amplicons were assembled into pAGM9121 [2] (Addgene ID #51833) using a GG cut-ligation reaction via BpiI.

pFH39: The rice *U3* promoter was amplified in fragments from the vector pPZP-FnCas12a-OsOpt [8] using primers FH27/FH30 and FH278/FH29. Both amplicons were assembled into pAGM9121 [2] (Addgene ID #51833) using a GG cut-ligation reaction via BpiI.

**Level 0 gRNA/crRNA backbone modules**

The vectors pFH85-pFH90 were synthesised by Twist Bioscience (San Francisco, USA).

The vectors pFH99-pFH104 were created by changing the GG overhang of pFH85-pFH90 via the Q5® Site-directed Mutagenesis Kit (New England Biolabs) using primers FH411/FH412 (on pFH85, pFH86, pFH87, pFH88) or primers FH413/FH414 (pFH89) or primers FH414/FH415 (pFH90) according to the supplier’s manual.

**Level 0 tRNA-sgRNA modules**

The modules pFH113 and pAK002-pAK012 were synthesised by GenScript (Piscataway, USA).

pFH49, pFH50, pFH51, pFH71: These modules were created by changing the GG overhang of pFH113 via the Q5® Site-directed Mutagenesis Kit (New England Biolabs) using primers FH303/FH304 (for pFH49), FH304/FH305 (for pFH50), FH304/FH306 (for pFH51) or primers FH304/FH323 (for pFH71) according to the supplier’s manual.

pFH72, pFH73, pFH74, pFH75: These modules were created by changing the GG overhang of pAK007 via the Q5® Site-directed Mutagenesis Kit (New England Biolabs) using primers FH303/FH304 (for pFH72), FH304/FH305 (for pFH73), FH304/FH306 (for pFH74) or primers FH304/FH323 (for pFH75) according to the supplier’s manual.

**Endlinkers**

The endlinker modules pAK-EL-01 – pAK-EL-05 were synthesised by GenScript (Piscataway, USA).

**Level 1 constructs**

pFH23, pFH58, pFH59, pFH60, pFH61, pFH62, pFH63, pFH64, pFH65, pFH66, pFH67, pFH68, pFH69, pFH70, pFH80, pFH81, pFH82, pFH84, pFH93, pFH109:

The respective level 0 nuclease constructs (pFH13, pFH14, pFH15, pFH16, pFH17, pFH18, pFH19, pFH20, pFH21, pFH32, pFH45, pFH55, pFH22, pFH24, pFH25, pFH76, pFH77, pFH78, pFH79, pFH92, pFH91) were assembled with the level 0 *ZmUBI* promoter-5’UTR level 0 construct (pICSL12009) [5] and a *NOS* 3’UTR-Terminator level 0 construct (pICH41421) [24] into the level 1 position 2 acceptor pICH47742 [2] (Addgene ID #48001) using a GG cut-ligation reaction via BsaI-HF®v2.

pFH52: An Arabidopsis codon optimised Cas9 (pICSL90016) [25] was assembled with a 2x35S promoter-5’UTR (pICH51288) [24] and a *NOS* 3’UTR-Terminator (pICH41421) [24] into the level 1 position 2 acceptor pICH47742 [2] (Addgene ID #48001) using a GG cut-ligation reaction via BsaI-HF®v2.

pFH53: An Arabidopsis codon optimised Cas9 (pICSL90016) [25] was assembled with a 2x*35S* promoter-5’UTR (pICH51288) [24] and a pea *3A* 3’UTR-Terminator (pFH44) into the level 1 position 2 acceptor pICH47742 [2] (Addgene ID #48001) using a GG cut-ligation reaction via BsaI-HF®v2.

pFH54: An Arabidopsis codon optimised Cas9 (pICSL90016) [25] was assembled with an *Arabidopsis* *UBIQUITIN10* promoter-5’UTR (pICSL12015) [25] and a pea *3A* 3’UTR-Terminator (pFH44) into the level 1 position 2 acceptor pICH47742 [2] (Addgene ID #48001) using a GG cut-ligation reaction via BsaI-HF®v2.

pFH114: The ZmUBIp::*BAR*:NOSt cassette was amplified using the vector pRRES1.111 [26] with FH443/FH444 primers and the PCR product was assembled into the level 1 position 1 acceptor pICH47732 [2] (Addgene ID #48000) using a GG cut-ligation reaction via BsaI-HF®v2.

pFH94 (tRNA-sgRNA array level 1 construct used for the protoplast assay): For integration of the 20 bp sgRNA target sequences into their respective level 0 tRNA-sgRNA subcloning vectors, primers (Sigma Aldrich) containing the 20 bp targets plus respective overhangs (TGCA for forward primers/AAAC for reverse primers) were adjusted to the concentration of 10 µM and mixed in the 1:1 ratio. The annealed primers were then cut-ligated into their respective tRNA-sgRNA level 0 vectors using BpiI (FH360/FH361 into pFH113, FH362/FH363 into pAK002, FH364/FH365 into pAK003, FH366/FH367 into pAK004, FH368/FH369 into pAK005 and FH370/FH371 into pAK006). The generated level 0 tRNA-sgRNA modules were then assembled into a transcription unit carrying the wheat U3 promoter (pFH31) into the level 1 backbone pICH47751 [2] (Addgene ID #48002) using BsaI-HF®v2.
